# OptoDyCE: Automated System for High-Throughput All-Optical Dynamic Cardiac Electrophysiology

**DOI:** 10.1101/023572

**Authors:** Aleksandra Klimas, Jinzhu Yu, Christina M. Ambrosi, John C. Williams, Harold Bien, Emilia Entcheva

**Affiliations:** Department of Biomedical Engineering, Stony Brook University, Stony Brook, NY, USA

## Abstract

The improvement of preclinical cardiotoxicity testing, the discovery of new ion-channel-targeted drugs, and the phenotyping and use of stem-cell-derived cardiomyocytes and other biologics all necessitate high-throughput (HT), cellular-level electrophysiological interrogation tools. Optical techniques for actuation and sensing provide instant parallelism, enabling contactless dynamic HT testing of cells and small-tissue constructs, not affordable by other means. Here, we consider, computationally and experimentally, the limits of all-optical electrophysiology when applied to drug testing, then implement and validate OptoDyCE, a fully automated system for all-optical cardiac electrophysiology. We validate optical actuation by virally introducing optogenetic drivers in (rat and human) cardiomyocytes or through the modular use of dedicated light-sensitive somatic “spark” cells. We show that this automated all-optical approach provides high-throughput means of cellular interrogation, i.e. allows for dynamic testing of >600 multicellular samples or compounds per hour, and yields high-content information about the action of a drug over time, space and doses.

---

The development of new drugs is a lengthy, inefficient process;^1^ in the United States, <0.05% of all compounds undergoing preclinical tests become marketed drugs, and <30% of compounds evaluated in expensive clinical trials make it to market.^2^ Perhaps most costly, and with greatest negative societal impact, is the withdrawal of drugs from the market after they have been approved. In the last two decades, over 30% of such withdrawals have been due to cardiac toxicity (pro-arrhythmic effects mediated by cardiac ion channels).^2^ International regulatory agreements mandate the testing of all new drugs for cardiac liability, including drug-induced long QT interval (LQT) and risk for development of life-threatening arrhythmias, such as Torsade de Pointes (TdP).^3^ This mandated preclinical testing specifically concerns a drug’s action as a potential blocker of the hERG K^+^ channel, which supplies one of the main repolarizing currents in cardiomyocytes. However, it has recently been recognized that a drug’s pro-arrhythmic effect, i.e. “torsadogenicity”, is often shaped by its action on multiple ion channels;^3-7^ therefore, an integrative (cell-level or multicellular) view is essential, and current regulations need to be revisited (see Supplementary Fig. 1 about the CiPA initiative, Comprehensive In Vitro Pro-arrhythmia Assay^3^). Computational efforts are underway^5,7^ to integrate multi-channel data obtained in recombinant expression systems to predict the action of a drug on the human cardiac action potential (see Supplementary Fig. 1). Recently, great strides in the optimization and production-scaling of human patient-derived cardiomyocytes (induced pluripotent stem cell derived, iPSC-CMs) promise to provide an alternative, more direct and relevant experimental testbed for cardiotoxicity screening.^8, 9^ However, there are currently no high-throughput solutions (ability to screen >10,000 compounds a day) for performing robust cardiomyocyte electrophysiological testing.

Classic electrophysiology involves physical contact and therefore is inherently very low throughput (manual). New technical developments towards increased throughput^10, 11^ include the automated planar patch, IonWorks by Molecular Devices, at the single-channel level; the Fluorometric Imaging Plate Reader (FLIPR) by Molecular Devices; Multichannel Electrode Arrays (MEAs) by Axion Biosystems; impedance-based assays with xCELLigence by Acea Biosciences; and the kinetic plate reader FDSS/μCell by Hamamatsu for cellular measurements (see **Supplementary Table 1** for a detailed comparison). The following limitations of these systems motivate the need for further developments towards HT cell-level electrophysiology: 1) requirements for contact in IonWorks, MEAs, or xCELLigence prevent scaling to the HT-level – a non-contact modality is a must; 2) lack of electrophysiologically-relevant fast readout (e.g. optical sensing in FLIPR makes it highly parallel, but still unable to track fast action potentials); 3) inability for dynamic actuation and frequency-response testing (e.g. FLIPR), which is quite relevant in drug-induced cardiotoxicity;^12^ 4) cell-type restrictions: more phenotypic outputs, such as iPSC-CMs or primary CMs as testbeds, are desirable instead of the currently-employed recombinant expression systems, but handling limitations present challenges (e.g. in IonWorks, a proper seal can only be formed with “well-behaved” cell lines^10^); 5) none of the current methods can characterize tissue-level/multicellular effects, even though arrhythmias are inherently spatiotemporal phenomena.

An all-optical electrophysiology approach^13-15^ can address these limitations and facilitate HT-level testing through built-in parallelism. New fast optogenetic tools for actuation^16-18^ and sensing^15, 19, 20^ offer attractive solutions for manipulating and observing multiple cells optically, but their limitations need to be considered carefully in the context of drug screening (Fig. 1a-g). Both actuators and sensors contain essential elements of ion channel proteins, making them susceptible to the drugs being tested. However, the action of a fast optogenetic actuator, e.g. Channelrhodopsin-2 (ChR2), in cardiomyocytes can be viewed as “time-detached” from the electrophysiological (EP) response, Fig. 1a, hence mostly benign. Computationally, we show that for brief light pulses, even dramatic drug effects on the ChR2 current amplitude and/or kinetics are practically inconsequential for the optically-triggered action potentials (APs) and calcium transients (CTs), if light irradiances are maintained at supra-threshold levels (Fig. 1b-f, details in **Supplement**). In contrast, an optogenetic sensor, e.g. VSFP2.3,^21^ is fully “temporally-convolved” with the EP response (Fig. 1a), and even a mild drug action on the sensor can profoundly alter the EP readout (see computational predictions, Fig. 1g); the same applies to other voltage^15, 20^ or calcium (GCaMP) optogenetic sensors, regardless of their kinetics. While great for long-term monitoring,^20^ channel-based optogenetic sensors may not be ideal for acute drug-testing applications due to such potential direct interference; instead, classic synthetic optical dyes (for voltage and calcium) may be more suitable, as they are already used in industrial applications.

**Figure 1.**
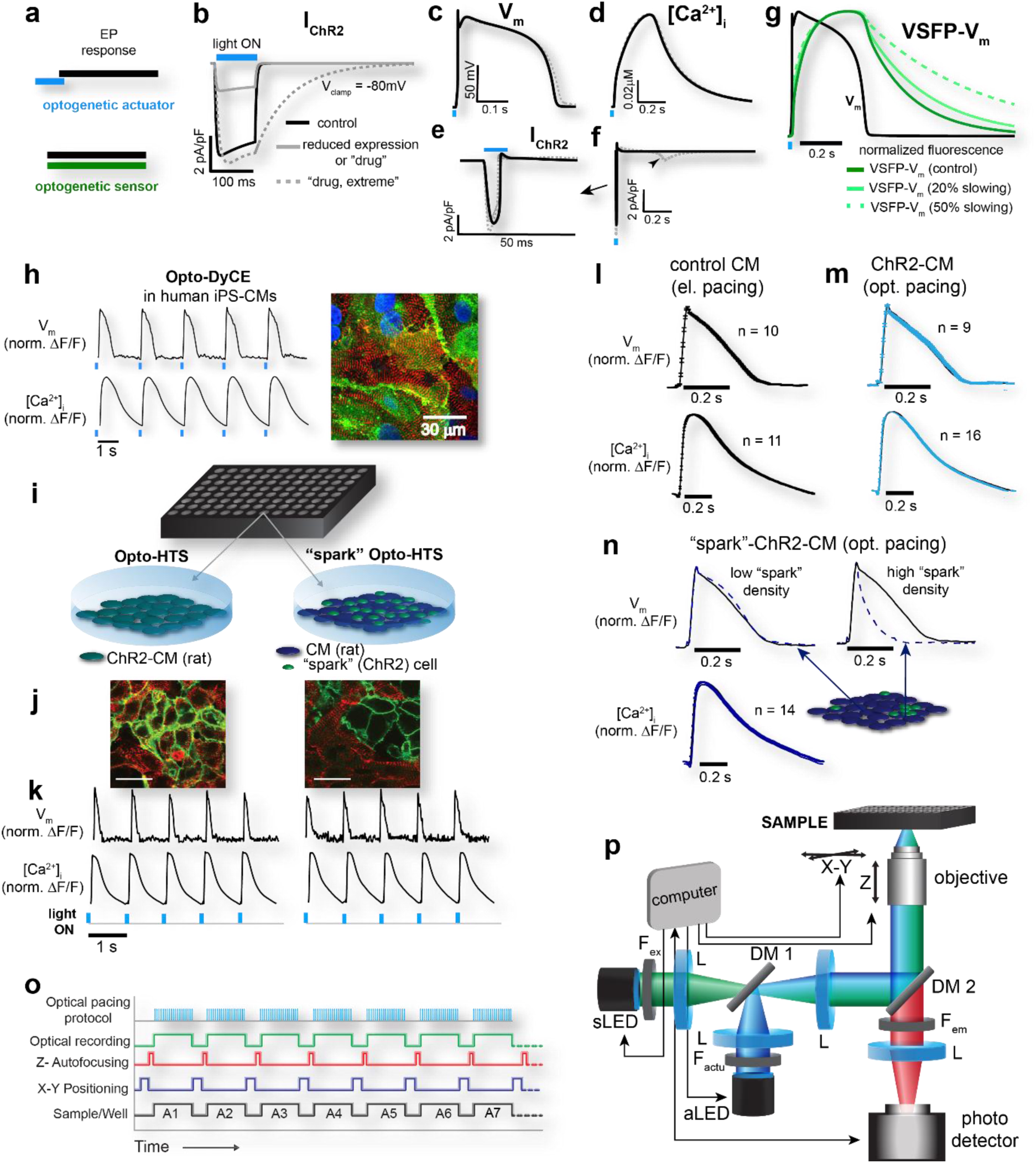
Feasibility, implementation strategies and validation of high-throughput all-optical electrophysiology. **(a-g)** Considerations and computational analysis of drug effects on optogenetic actuators (ChR2) versus optogenetic sensors (VSFP2.3) in human ventricular myocytes. A fast optogenetic actuator is “time-detached” from the electrophysiological (EP) response **(a)**, and therefore a hypothetical drug action that affects ChR2 current amplitude and/or kinetics **(b)** has minimal effect on the optically-triggered action potentials (APs) **(c)** and calcium transients (CTs) **(d)** if light irradiances are adjusted to supra-threshold currents **(e)**. Even extreme drug interference with ChR2 off-kinetics results in minor (5%) APD prolongation **(c)** due to re-activation of inward ChR2 current during repolarization **(f)**. In contrast, an optogenetic sensor, e.g. VSFP2.3, is fully temporally-convolved with the EP response **(a)**, and even a mild drug action on the sensor can profoundly influence the EP readout **(g)**. **(h-p)** High-throughput implementation of OptoDyCE. Human ChR2-iPSC-CMs **(h)** as well as rat ChR2-CM (OptoHTS) and “spark”-ChR2-CMs (sOptoHTS) **(i-k)** are optogenetically transformed to respond to optical stimulation. ChR2 expression by eYFP reporter (green), α-actinin staining (red) and DAPI nuclear stain (blue) are shown **(h, j)**. Optical pacing reliably triggers V_m_ and [Ca^2+^]_i_ signals, measured optically **(h, k).** Validation of OptoHTS comes from identical AP and CT morphology for electrically-paced CM-controls (non-transduced) and optically-paced ChR2-CMs **(l-m)**; in sOptoHTS, high “spark” cell density can lead to APD shortening compared to control CM (**n**, Supplementary Fig. 2) without much effect on CT morphology **(n).** We demonstrate a fully automated high-throughput version of OptoDyCE in 96-well format using a custom-built optical setup and an automation protocol **(o-p)**, details in **Supplement**.

Here, we present *OptoDyCE*, the first fully automated platform for all-optical dynamic interrogation of cardiomyocyte electrophysiology, including human iPSC-CMs (Fig. 1h, Supplementary Fig. 2), with applicability to drug testing. We demonstrate the HT capabilities of OptoDyCE using multicellular samples in 96-well format by combining optogenetic actuation (via ChR2) with simultaneous optical sensing of voltage or intracellular calcium by synthetic red-shifted dyes (di-4-ANBDQBS and Rhod-4AM, respectively), illustrated in Fig. 1h-p. Contactless optical pacing reliably triggers voltage (V_m_) and calcium ([Ca^2+^]_i_) signals (Fig. 1h, k). We impart the ability for optical pacing via one of two quick and efficient transduction methods (within 48 hours prior to experimentation) to yield: 1) *OptoHTS*: using direct adenoviral gene delivery (in human, ChR2-hiPSC-CM, or neonatal rat ventricular cardiomyocytes, ChR2-CM);^13, 22, 23^ or 2) *sOptoHTS*: “sprinkling” of dedicated light-sensitive (“spark”) cells on top of samples of non-transduced cardiomyocytes, a version of our “tandem-cell-unit” concept,^24^ (Fig. 1i-k, see **Supplement**).

**Figure 2:**
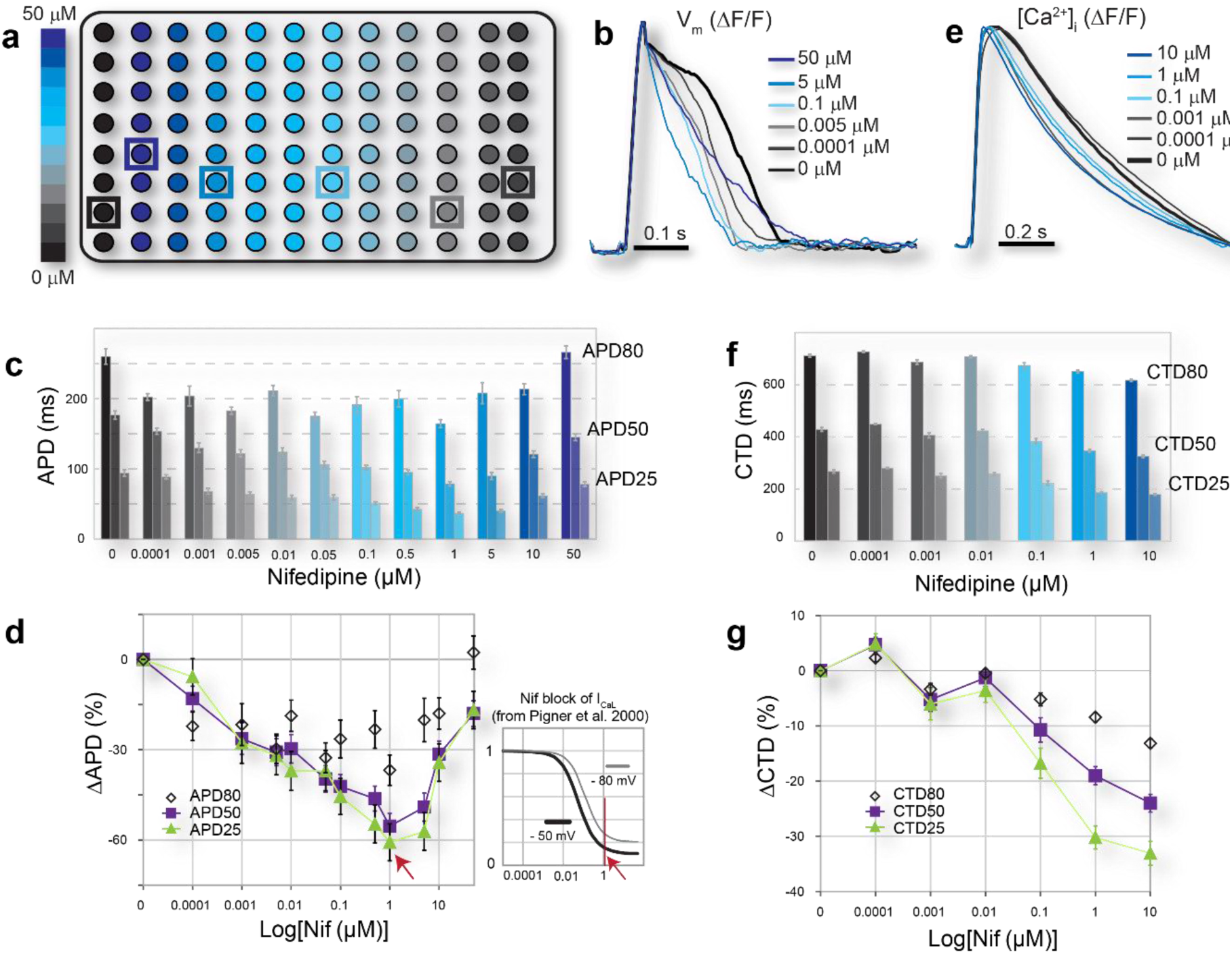
Demonstration of OptoHTS for high-throughput dose-response drug testing. Nifedipine, an L-type Ca^2+^ channel (I_CaL_) blocker, is applied at 12-concentration graded dosing (0 – 50 μM) to ChR2-CMs in 96-well plates **(a)**. Optical recordings of multiple voltage **(b-d)** or calcium events **(e-g)** are obtained during optical pacing at 1Hz, screening the full plate in under 10 minutes (see also Supplementary Fig. 3). Example averaged traces **(b, e)** and quantitative results for APD and CTD **(c-d, f-g)** are shown. Expected APD **(b-d)** and CTD **(e-g)** shortening, especially at the APD25/CTD25 and APD50/CTD50 levels, occurred due to nifedipine blocking the inward L-type calcium current. Maximum APD shortening is observed at around 1 μM, consistent with maximum block of I_CaL_ reached at that concentration^27^ **(d inset)**. Beyond 1 μM, indirect (voltage-mediated) or non-specific action on other ion channels partially counters the block of inward Ca^2+^ current and can reduce or eliminate the APD shortening **(d).** Nifedipine appears to monotonically shorten CTD up to 10 μM **(f,g)**. Data are presented as mean±SEM, and each well is considered an independent sample, represented by a spatially-averaged trace.

We validate OptoHTS by comparing AP and CT morphology of optically-stimulated ChR2-CM samples and electrically-paced non-transduced CM samples, confirming optogenetic pacing is a suitable alternative to electrical stimulation for drug testing purposes (Fig. 1l-m), as predicted computationally.^22, 25^ The sOptoHTS provides a more attractive modular method of light sensitization, as a bank of generic “spark” cells (light-sensitized somatic cells) can be used in conjunction with a variety of non-modified experimental cardiac systems. However, caution should be applied regarding the geometry of “spark” cell distribution, since loading effects of higher “spark”-cell concentrations can locally shorten the AP (Fig. 1n, Supplementary Fig. 3), having minimal effects on CT morphology. Proper “spark”-cell delivery (for example, a localized pacing site) can easily address the issue.

**Figure 3:**
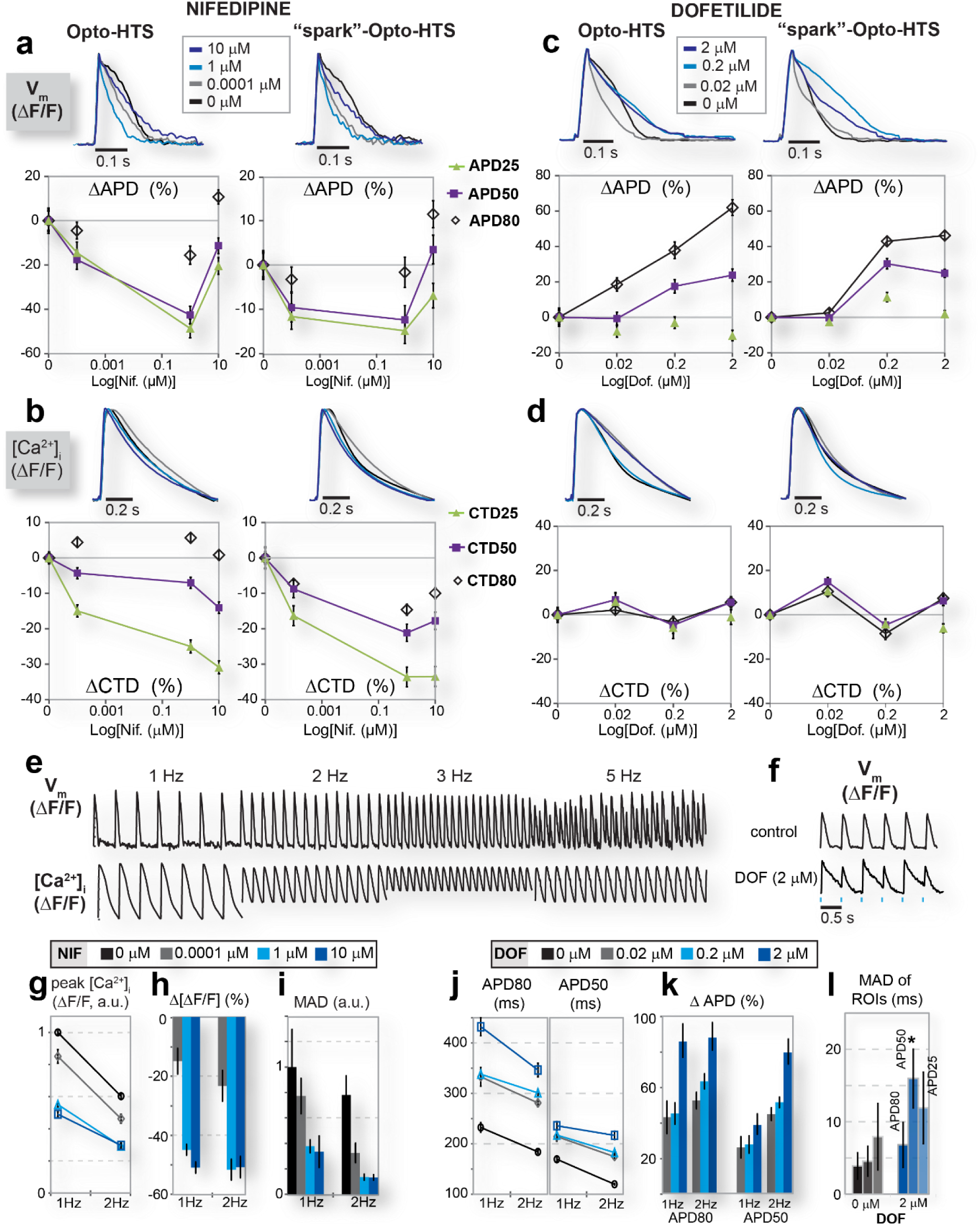
Comparison of OptoHTS versus “spark”-OptoHTS for dynamic functional drug testing. **(a-d)** OptoHTS (left) and sOptoHTS (right) provide qualitatively and quantitatively similar results for measured effects on APD **(a, c)** and CTD **(b, d)** for both nifedipine **(a,b)** and dofetilide, a blocker of the rapid delayed rectifier, I_Kr_ **(c,d)**. The ability to perform dynamic pacing **(e-l)** provides a means of studying pacing-induced V_m_ and Ca^2+^ restitution and instabilities **(e)** or drug-induced instabilities, i.e. 2 μM dofetilide leading to voltage alternans at relatively low pacing frequency (2Hz) **(f)**. High-content dynamic information is obtained from a single data run **(g-l)**. For example, restitution and temporal or spatial variability (quantified by median absolute deviation (MAD)) are shown as function of both drug dose and pacing frequency for peak calcium in the presence of nifedipine **(g-i)** and for APD in the presence of dofetilide **(j,k)**. Nifedipine action on peak calcium (% change) is dose-dependent but frequency-independent **(g,h).** Nifedipine appears to reduce temporal variability of peak calcium (assessed by MAD), and this reduction is augmented by higher-frequency pacing **(i)**. Dofetilide shows enhanced action on APD50 at higher frequency (opposite to reverse-use dependence) **(j,k)**. Spatial variation as a function of drug dose can also be assessed by analysing multiple regions of interest (ROIs) within the same well **(l,** see also Supplementary Fig. 3). Dofetilide at 2 μM seems to increase spatial variability in APD, i.e. increase dispersion of repolarization, compared to control during 1Hz pacing (p<0.05 for APD50). Five to 16 multicellular samples were used for each condition and each data point in the panels above.

A fully automated high-throughput version of OptoDyCE in 96-well format is demonstrated here using an optical setup custom-built around an inverted microscope, an automation protocol, and custom-developed software for semi-automated analysis (Fig. 1o-p, Supplementary Fig. 4-5, details in **Supplement**). The simplicity of our implementation makes it easy to adopt; compared to a prior report on all-optical electrophysiology in neurons,^15^ we use low-power LED light sources and standard equipment.

To our knowledge, this is the first scalable automated all-optical platform for cardiac electrophysiology that can meet the HT standard (see considerations in Supplementary Fig. 4). With robotic dispensing, the 96-well format can be instantly upgraded to 384-well or other standard plate formats, with very simple reprogramming. The current implementation has built-in parallelism within a well, interrogating hundreds of cells simultaneously (Supplementary Fig. 4), but relies on serial traversing of the wells; a *macroscopic* version (see^26^ or in FDSS/μCell) of OptoDyCE with larger field-of-view can further increase throughput by order(s) of magnitude.

Even in the current proof-of-concept implementation of OptoDyCE, dynamic drug-dose testing can be performed on a 96-well platform (over 30,000 single-cell readouts, multi-beat pacing protocol) *in less than 10 minutes* (Fig. 2a, Supplementary Fig. 4). As an illustration, nifedipine, a class-IV antiarrhythmic agent, was applied in 12 doses (0 to 50μM) to 96 ChR2-CM samples (Fig. 2a). Using optical pacing at 1Hz, we quantified the dose-response to nifedipine in terms of AP duration (APD) and CT duration (CTD) (Fig. 2b-g). Expected APD shortening (Fig. 2b-d) and CTD shortening (Fig. 2e-g) was observed, especially at the plateau phase (APD25/CTD25 and APD50/CTD50), due to nifedipine blocking the inward L-type calcium current, I_CaL_. Nifedipine caused CTD to monotonically decrease up to 10 μM (Fig. 2f,g). In contrast, after maximum APD shortening at around 1 μM, corresponding to maximum block of I_CaL_ reached at that concentration^27^ (Fig. 2d inset), the APD response to nifedipine reversed its direction, as seen clinically.^6^ This was likely due to indirect (voltage-mediated) or non-specific action on other ion channels, partially countering the I_CaL_ block (Fig. 2d). Note that the benefits of our *in vitro* HT platform are in the ability to quickly and finely probe many concentrations and to help determine the “therapeutic window”, i.e. the window for which a drug is both effective (has the desired action) and safe. Clinically, the drug-metabolizing action of the cytochrome P450 enzymes can amplify or suppress the effect of a drug which can can result in a lower or higher *apparent drug dose* (as in some failed drugs^2^); our data can be used to judge the “room for error” in the therapeutic window for a drug.

Seeking validation for sOptoHTS, we further compared the dose-dependent effects of nifedipine and of dofetilide using the two methods, OptoHTS vs. sOptoHTS (Fig. 3). Dofetilide, a class-III anti-arrhythmic agent and intended hERG channel blocker, has a known risk for drug-induced LQT and TdP due to its APD-prolonging action.^6^ sOptoHTS was able to successfully track the drug-dose dependent effects on APD and CTD, similar to OptoHTS (Fig. 3a-d). With proper tuning of the “spark” cell distribution, this simple and truly modular approach provided by sOptoHTS can be reliably applied to HT electrophysiological drug testing.

Electrophysiological responses are frequency-dependent, therefore passive observation of spontaneous activity^20^ is generally insufficient in drug testing. Unlike most currently-employed systems (**Supplementary Table 1**), our platform allows for active dynamic interrogation, such as robust pacing protocols that can reveal V_m_ and [Ca^2+^]_i_ frequency response (restitution) and temporal instabilities (Fig. 3e). For example, a consistent generation of voltage instabilities known as alternans can be captured at 2Hz optical pacing in the presence of 2 μM dofetilide due to drug-induced APD prolongation (Fig. 3f). Restitution and temporal or spatial variability (assessed by median absolute deviation (MAD), see **Supplement**) can be quantified as function of drug dose (Fig. 3g-l). These are directly relevant to the “torsadogenecity” of a drug, providing a much more complete assessment than traditional (single-channel block) testing or current state-of-the-art assays (**Supplementary Table 1**). Our dynamic testing data reveal that nifedipine action on peak calcium (% change) is dose-dependent (p<0.05) but frequency-independent (Fig. 3g-h). Furthermore, nifedipine reduces temporal variability of peak calcium (assessed by MAD), and this reduction is augmented by higher-frequency pacing (Fig. 3i). For dofetilide, we found enhanced relative APD50 prolongation at higher frequency, which is opposite to purported reverse-use dependence (Fig. 3j-k). Furthermore, because of the ability to study multicellular samples, spatial variability can be quantified as a function of drug dose by analysing individual cells or regions of interest within the same sample/well (Fig. 3l, see also Supplementary Fig. 4). For example, we found that dofetilide at 2 μM increases spatial variability in APD (i.e. increases dispersion of repolarization -- a known pro-arrhythmic factor), compared to control during 1Hz pacing (p<0.05 for APD50). Most of the electrophysiological results presented in Figs. 2-3 can be corroborated by published work, albeit certainly not in an all-encompassing study. We chose to conduct our demonstration with these known clinically-used drugs so that the emphasis of our study can be the illustration of usability and the potential impact of our methodology rather than the electrophysiological insights per se.

These results also illustrate the high-content data that can be obtained with OptoDyCE, that allows the quantification of a drug’s pro-arrhythmic risk in a much more comprehensive way than with any of the current platforms. The contactless nature of interrogation in OptoDyCE makes it especially versatile and applicable to not only single cells and cell monolayers, but also to small-thickness 3D engineered tissues (within the optical penetration depth). By offering a currently missing option for automated HT cardiomyocyte electrophysiology, OptoDyCE can profoundly impact developments concerning human iPSC-CMs^8, 9^ (see Supplementary Fig. 1) by allowing for *combinatorial optimization* of factors involved in cell maturation, phenotype selection, and tissue engineering. In turn, the utilization of these new optimized human (potentially patient-specific) experimental models, in conjunction with our HT testing platform, has the potential to dramatically improve pre-clinical drug testing, reduce cost, reduce animal use, and increase a therapy’s likelihood of clinical success.

## Acknowledgements

We thank Nathaniel Hobert for help with data analysis. This work was supported by grants to E.E. from the NIH-NHLBI HL111649 and from the NSF-Biophotonics division 1511353, as well as a NYSTEM grant C026716 to the Stony Brook Stem Cell Centre.

### Author Contributions

E.E. conceived the project, designed the experiments and provided guidance. A.K., J.Y. and C.M.A. assisted in the experimental design. A.K. built the optical system and designed the automation routines. H.B. and A.K. wrote the software to collect and analyse the data. C.M.A. developed and tested the optogenetic tools used in the project. A.K., J.Y. and C.M.A. performed the experiments. J.C.W. and E.E. performed the computational analysis. A.K. and E.E. wrote the manuscript. All authors were involved in discussion of the results and revision of the manuscript.

Supplemental Information

*for Klimas A et al., OptoDyCE: Automated System for High-Throughput All-Optical Dynamic Cardiac Electrophysiology*

## Supplemental Methods

***(Detailed Supplemental Methods not included in this document)***

## Supplemental Figures

**Supplemental Figure 1:**
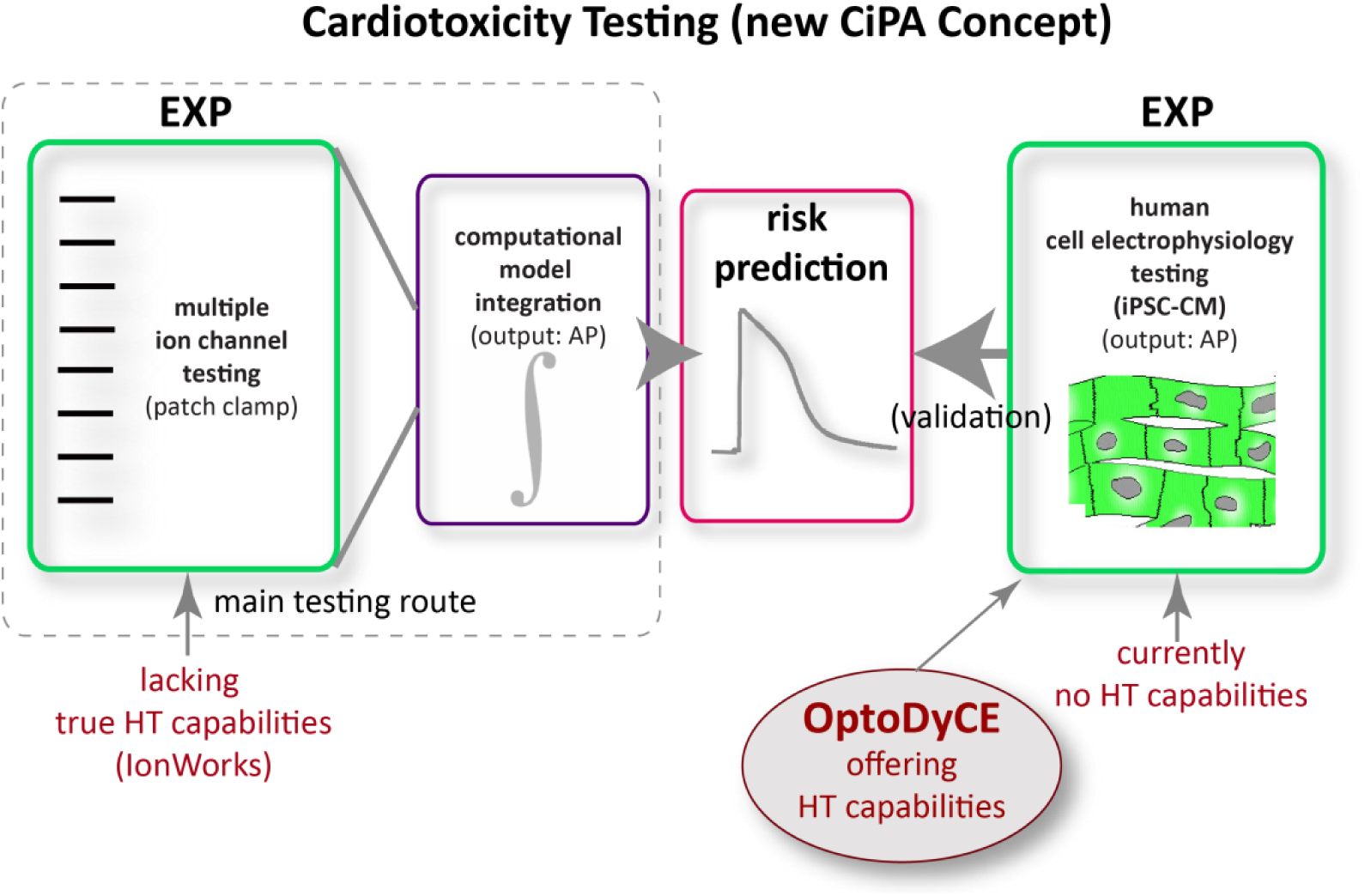
Cardiotoxicity testing: the new CiPA^11^ (Comprehensive In Vitro Pro-arrhythmia Assay) concept and a role for OptoDyCE. The new CiPA concept aims to help change/improve current ICH* regulations for pre-clinical cardiotoxicity testing to avoid unnecessary drug attrition, reduce cost and improve drug development. The arrhythmia risk is to be derived from experimental data using multi-channel testing by manual or planar automated patch clamp in recombinant expression systems (see IonWorks in **Supplemental Table 1**), which are then integrated using computational tools to predict the overall action of a compound on human cell electrophysiology, i.e. on the action potential (AP) - this experimental step plus the computational step form the main testing route. The predictions (the effects of a compound on the human action potential) are to be validated using cell electrophysiology measurements in human cardiomyocytes (most likely, iPSC-CMs). Note that currently, both of the two experimental components in this scheme lack true high-throughput (HT) capabilities. Planar patch systems have evolved but do not pass the HT threshold; cardiomyocyte electrophysiology (AP measurements) currently cannot be performed in HT fashion. Our platform, **OptoDyCE, aims to bring HT capabilities to the cell-level testing in human cardiomyocytes**. This experimental approach (on the right) is more direct and can theoretically (pending maturation of the iPSC-CM technology) provide more relevant, and even patient-specific predictions, compared to the main testing route on the left (a comprehensive characterization of all ion channels is impossible, and hence the complex computational models operate in high level of uncertainty, thus providing only probabilistic predictions). Furthermore, OptoDyCE can provide additional cell- and multicellular readouts, e.g. intracellular calcium, cell coupling, which are very relevant to arrhythmia testing but cannot be derived by the approach on the left. Therefore, OptoDyCE can help further improve computational modeling as well.

**Supplemental Figure 2:**
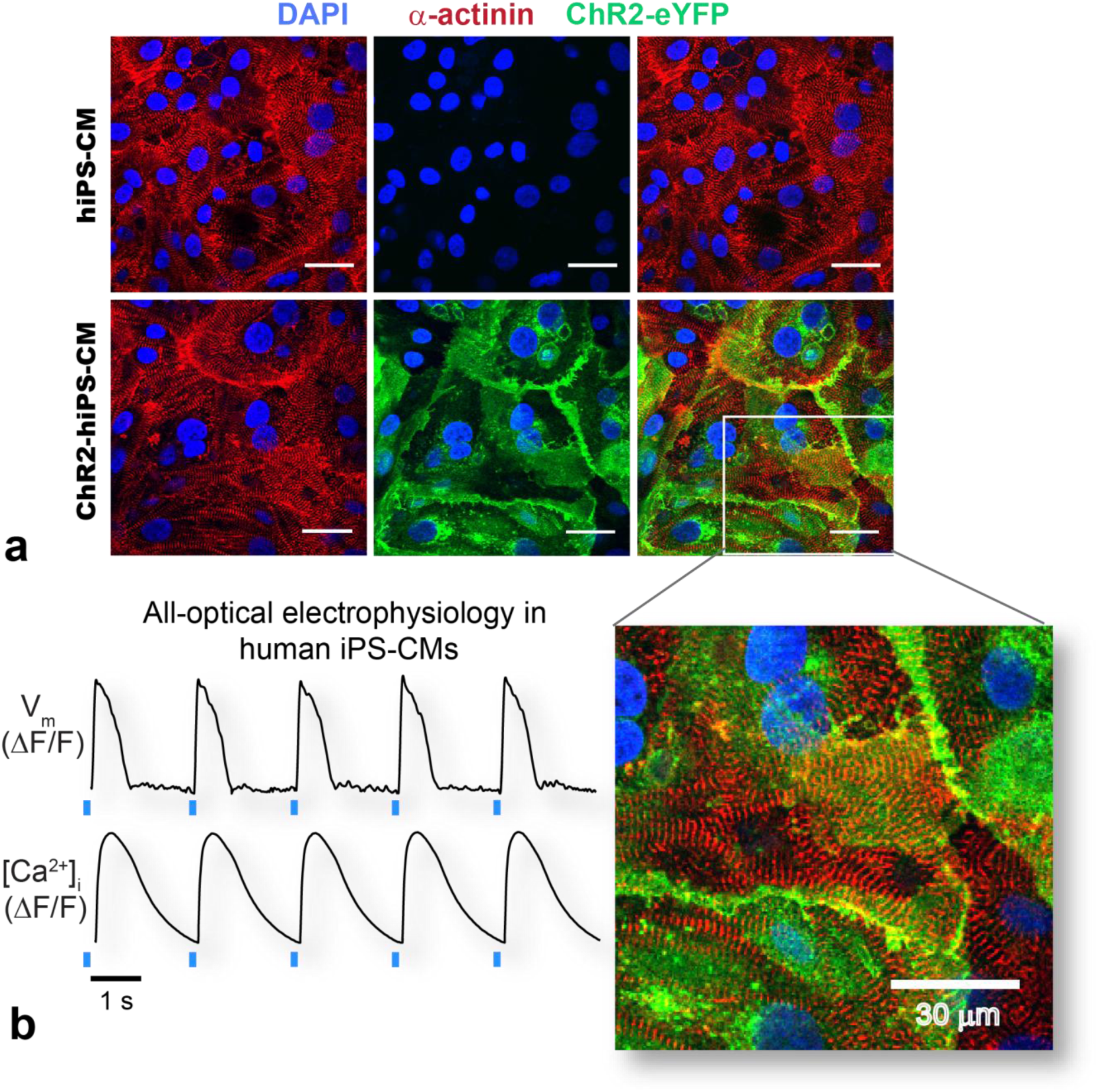
All-optical electrophysiology in human iPS-CMs (expanded Fig. 1h). **(a)** hiPS-CMs without (top) and with Ad-ChR2(H134R)-eYFP delivery at MOI 250 (bottom). Red fluorescence indicates α-actinin staining illustrating the cardiomyocyte-like properties of hiPS-CMs, blue indicates DAPI nuclear staining, and green fluorescence indicates the eYFP reporter of ChR2. Combination (left) of the α-actinin (right) and eYFP (center) channels indicate expression of ChR2 in the ChR2-hiPS-CMs. **(b)** Optical recording of V_m_ and [Ca^2+^]_i_ in optically paced ChR2-hiPS-CMs, used in an automated readout (96-well plate format).

**Supplemental Figure 3:**
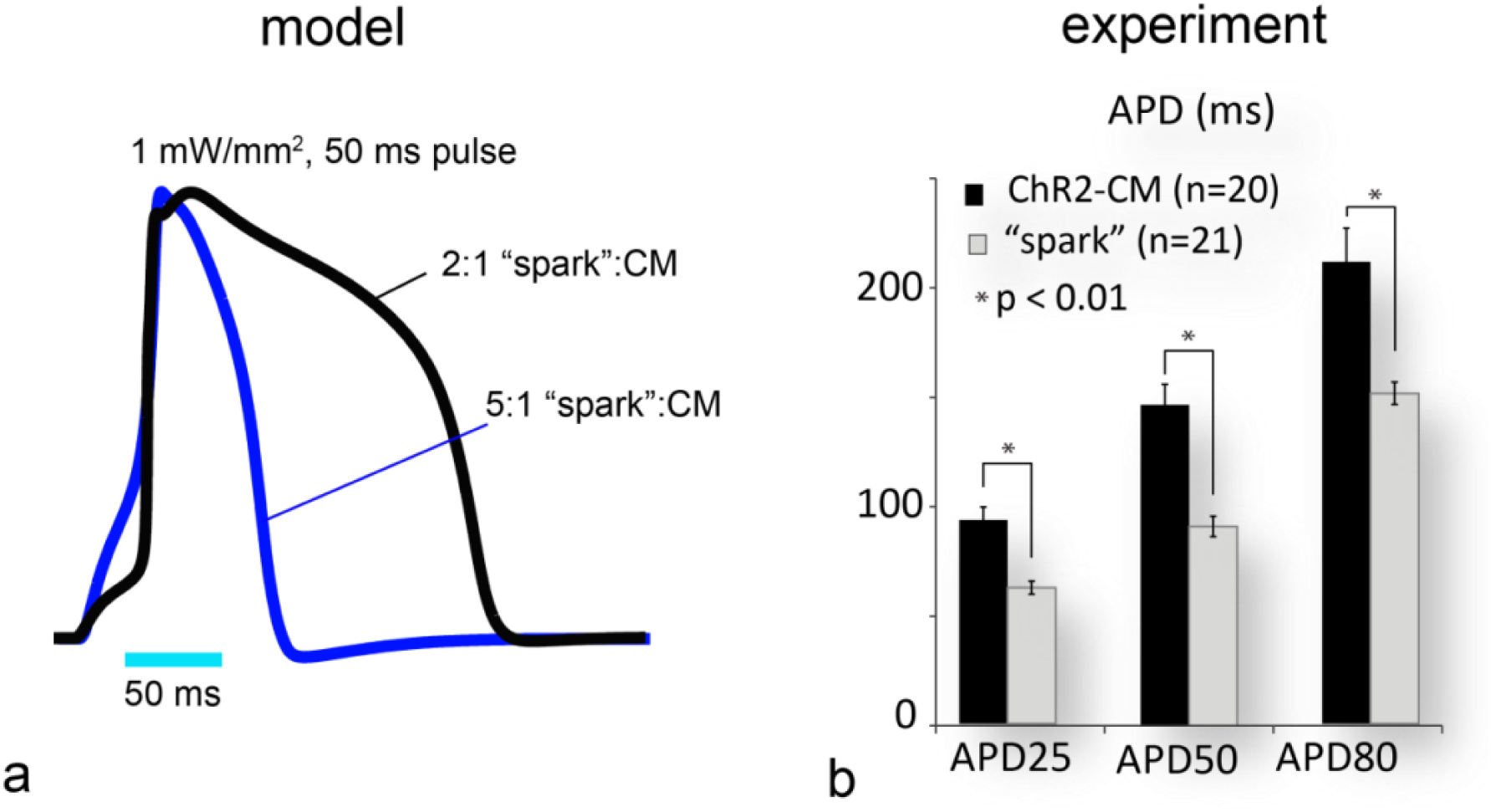
Effect of “spark” cell density on measured APD in CMs. **(a)** Shown is an example of human ventricular action potentials, scaled and normalized in amplitude (as would be measured by an optical method) for two cases of “spark”-driven excitation of CMs: when 2 “spark” cells were connected to a cardiomyocyte or when 5 “spark” cells were connected to a cardiomyocyte. The loading effect in the latter case resulted in APD shortening in the CM (see Fig. 1n). The computer model employed ChR2-expressing cardiac fibroblasts (not HEK cells) as “spark” cells (see *Computational Analysis*), but the effect is applicable to both cell types. **(b)** Comparison of the experimentally measured APD for optically paced ChR2-CMs and HEK-ChR2-CMs for the samples shown in Fig. 1m-n. For the cell density and the implementation here, with random sprinkling, there was overall APD shortening in the “spark”-driven (HEK-ChR2-CMs) compared to the ChR2-CMs (p < 0.01); to avoid APD shortening, the “spark” cells can easily be localized and can serve as optical pacemaking conduits without affecting the APD of the cardiomyocytes.

**Supplemental Figure 4:**
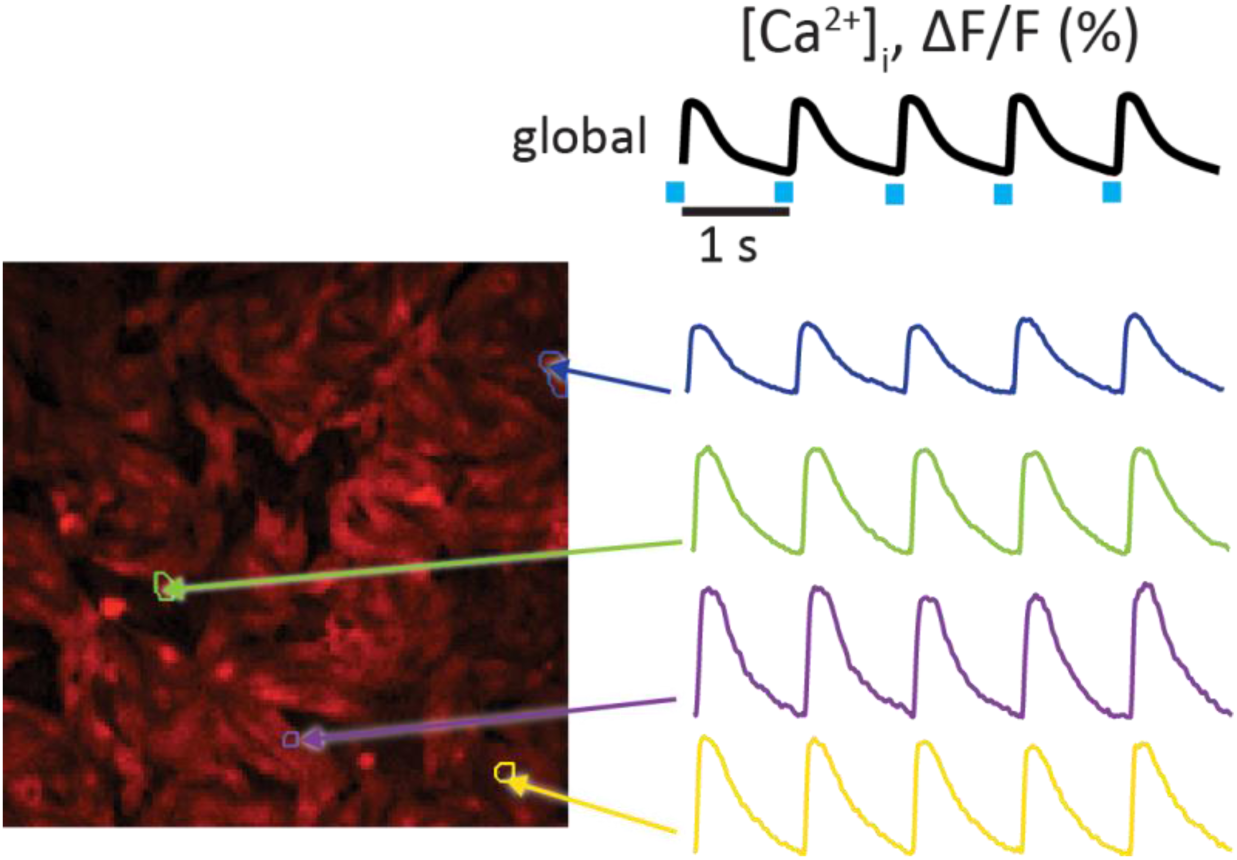
Parallelism of the OptoHTS system. In each well, optical recording was performed over multiple cells in parallel. Typical field of view (FOV) was 400 μm × 400 μm, resulting in about 200-400 parallel cell measurements per FOV (well), i.e. about **30,000 cell-level measurements per 96-well plate** at 20x magnification. For a dynamic pacing protocol, using **multi-beat pacing** (6 second dwell time per well), this resulted in about **10 min/plate**, i.e. about **600 independent multi(>200)-cellular samples (or compounds) per hour** (with the possibility to reach **> 10,000 compounds per day**, which qualifies for HTS). Shown here are the global (space-integrated) calcium measurement for a well and traces from individual cell-level regions, as outlined. While most of the analysis presented here dealt with the global responses, the parallelism is built-in into our approach and can easily be utilized further (see Fig. 3l).

**Supplemental Figure 5:**
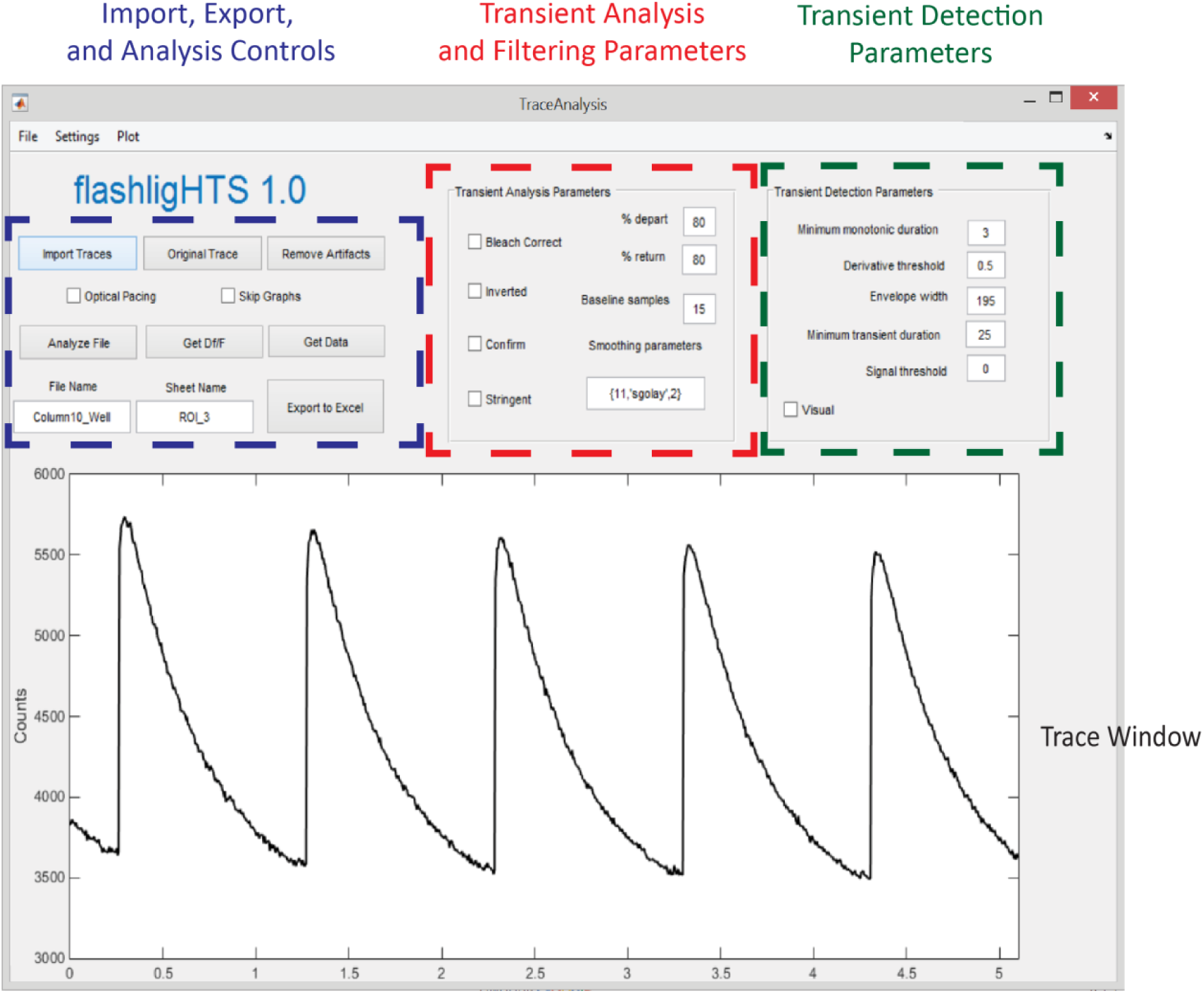
flashligHTS analysis software. A snapshot of the custom-developed automated analysis software is shown.

**Supplemental Figure 6:**
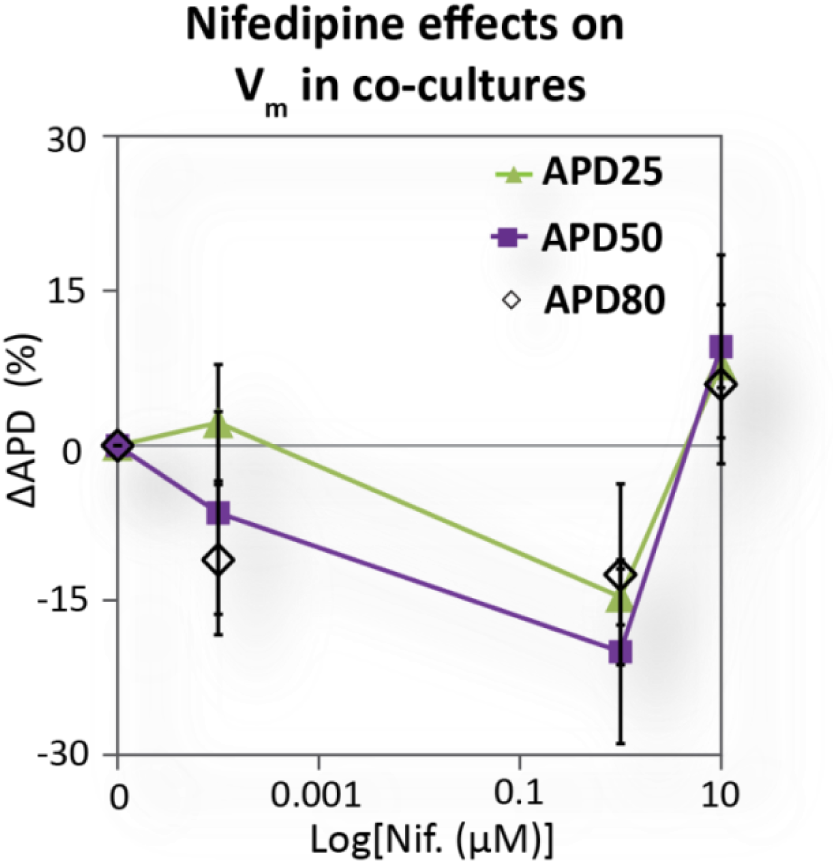
Drug action in standard TCU co-cultures. Shown is the effect of nifedipine on APD in standard co-cultures based on the TCU concept, when the “spark” cells are not “sprinkled” at a later point but rather mixed uniformly at the time of plating of the CMs. Comparing with Fig. 3a, all three cases qualitatively capture the action of nifedipine on APD, but the sOptoHTS with “sprinkling” is the simplest, most modular and attractive for industrial application.

**Supplemental Figure 7:**
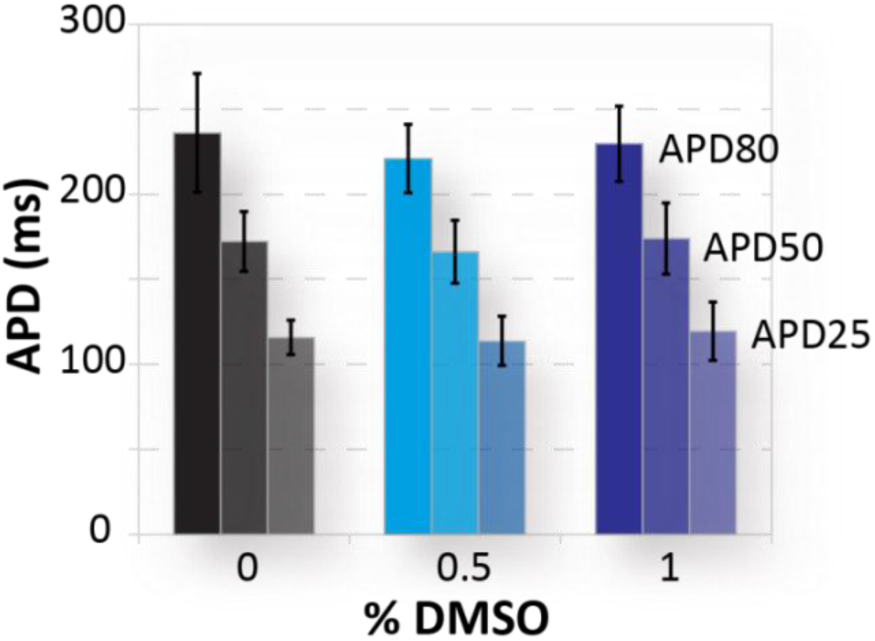
Effect of DMSO on APD. CM-ChR2 cells were dosed with 0, 0.5, and 1 % DMSO in Tyrode’s solution to assess the effect of DMSO on APD. Within the used concentrations to administer drugs and/or dyes (in all cases <1%), DMSO was not seen to affect the electrophysiological measurements.

**Supplemental Table 1.** State-of-the art industry-employed high-throughput approaches for electrophysiological measurements

| System | Description | PROS | CONS |
| --- | --- | --- | --- |
| **IonWorks**<br>*Molecular Devices*<br>12-17 | **Actuation:**<br>Electrical<br>**Sensing:**<br>Population patch clamp | • Closest to classic ion channel characterization<br>• Fast readout<br>• Dynamic stimulation available | • Limited throughput/expandability<br>• Limited spatial resolution<br>• Contact-requiring<br>• Not robust (needs “well behaved” cell lines; not applicable to any primary cells or mini-tissues)<br>• No calcium/contractility measurements<br>• High complexity; custom plates<br>• High cost |
| **Maestro Multichannel Electrode Array (MEAs)** *Axion Biosystems*<br>12, 18-21 | **Actuation:**<br>Electrical<br>**Sensing:** Local field potential | • Label-free<br>• Long-term recording possible<br>• Fast readout<br>• Dynamic stimulation available | • Limited throughput/expandability<br>• Limited spatial resolution<br>• Contact-requiring<br>• Not robust (not applicable to mini-tissues)<br>• No direct measurements of action potentials, calcium or contractility<br>• High complexity; custom plates |
| **xCELLigence**<br>*Acea Biosciences*<br>18, 19, 22, 23 | **Actuation:**<br>Electrical<br>**Sensing:**<br>Impedance | • Label-free<br>• Long-term recording possible | • Limited throughput/expandability<br>• Tracks only slow processes<br>• Limited spatial resolution<br>• Contact-requiring<br>• Not robust (not applicable to mini-tissues)<br>• No direct measurements of action potentials, calcium or dynamic contractions<br>• High complexity; custom plates<br>• High cost |
| **FLIPR**<br>*Molecular Devices*<br>13, 17, 24, 25 | **Actuation:**<br>Chemical<br>**Sensing:** Optical (fluorescence) | • Highly parallel<br>• Contactless<br>• Optical readout | • No dynamic stimulation<br>• Slow readout<br>• No spatial information<br>• Not robust (not applicable to mini-tissues)<br>• No direct measurements of action potentials or contractility<br>• High complexity; custom plates<br>• High cost |
| **FDSS/μCell**<br>*Hamamatsu*<br>24, 26 | **Actuation:**<br>Chemical (or electrical field)<br>**Sensing:** Optical (fluorescence) | • Highly parallel<br>• Contactless<br>• Optical readout<br>• Dynamic stimulation available | • Limited dynamic stimulation<br>• Relatively slow readout (typical < 5fps)<br>• No spatial information (low SNR)<br>• Not robust (not applicable to mini-tissues)<br>• No direct measurements of action potentials or contractility<br>• High complexity; custom plates<br>• High cost |

## Footnotes

* ICH - International Conference on Harmonization of Technical Requirements for Registration of Pharmaceuticals for Human Use – coordinated regulatory efforts by Europe, Japan and the United States concerning pharmaceutical products.

